# Microbial-induced calcium carbonate precipitation: An experimental toolbox for in situ and real-time investigation of micro-scale pH evolution

**DOI:** 10.1101/2020.04.15.042168

**Authors:** Jennifer Zehner, Anja Røyne, Alexander Wentzel, Pawel Sikorski

## Abstract

Concrete is the second most consumed product by humans, after water. However, the production of cement, which is used as a binding material in concrete, causes more than 5% of anthropogenic CO_2_ emissions and has therefore a significant contribution to climate change and global warming. Due to increasing environmental awareness and international climate goals, there is a need for emission-reduced materials, that can replace conventional concrete in certain applications. One path to produce a solid, concrete-like construction material is microbial-induced calcium carbonate precipitation (MICP). As a calcium source in MICP, crushed limestone, which mainly consists out of CaCO_3_, can be dissolved with acids, for example lactic acid. The pH evolution during crystallization and dissolution processes provides important information about kinetics of the reactions. However, previous research on MICP has mainly been focused on macro-scale pH evolution and on characterization of the finished material. To get a better understanding of MICP it is important to be able to follow also local pH changes in a sample. In this work we present a new method to study processes of MICP at micro-scale *in situ* and in real time. We present two different methods to monitor the pH changes during the precipitation process of CaCO_3_. In the first method, the average pHs of small sample volumes are measured in real time, and pH changes are subsequently correlated with processes in the sample by comparing to optical microscope results. The second method is introduced to follow local pH changes at a grain scale in *situ* and in real time. Furthermore, local pH changes during the dissolution of CaCO_3_ crystals are monitored. We demonstrate that these two methods are powerful tools to investigate pH changes for both MICP precipitation and CaCO_3_ dissolution for knowledge-based improvement of MICP-based material properties.

**Graphical TOC Entry:** 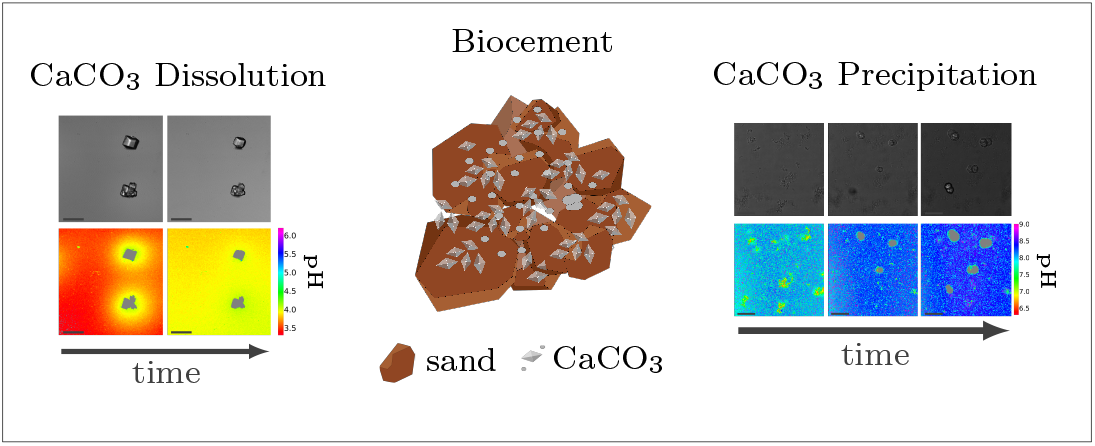

## Introduction

The concrete industry is a major contributor to human-made CO_2_ emissions. The production of cement as a binder in concrete for construction purposes accounts for more than 5% of all anthropogenic CO_2_ emissions^1^. Up to 65% of the CO_2_ created during cement production derives from the chemical decomposition of limestone (CaCO_3_) to lime (CaO). High temperatures are necessary for this decomposition process. The burning of fossil fuels for reaching high temperatures during the production procedure accounts for the remaining part of the CO_2_ emissions from cement production^2^.

Due to increasing awareness of the climate impact of anthropogenic CO_2_, there is a large interest in reducing the global CO_2_ emissions, including the carbon footprint of the construction industry. Approaches such as alternative raw materials, alternative fuels, and carbon capturing techniques have the potential of energy savings and CO_2_ reduction in cement production^3–5^. Another approach is to replace conventional concrete in some applications with a material based on biomineralization, so-called bio-cement. The production procedure of bio-cement utilizes bacteria to achieve biomineralization within a granular material to form a solid, concrete-like material. The most researched mechanism for producing bio-cement is microbially induced calcium carbonate precipitation (MICP)^6–9^. Ureolytic bacteria such as *Sporosarcina pasteurii* have been studied extensively in MICP experiments^10–12^. The chemical process of MICP is based on hydrolysis of urea^10,13^. The enzyme urease catalyzes the hydrolysis to ammonia and carbamate (Equation 1), which further hydrolyzes spontaneously to ammonia and carbonic acid (Equation 2):

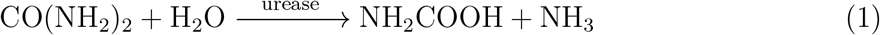

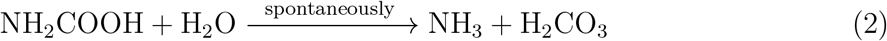

Ammonia and carbonic acid equilibrate further to form ammonium and bicarbonate (Equation 3, Equation 4):

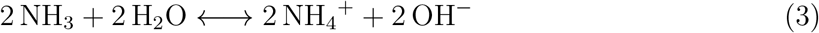

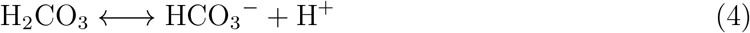

Resulting hydroxide ions lead to an increase in the pH and a shift in the bicarbonate equilibrium. In the presence of sufficient concentration of calcium ions this can result in CaCO_3_ precipitation:

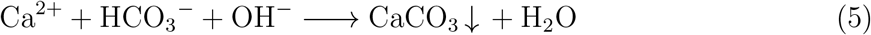

The MICP process is controlled by four key-factors: (1) availability of nucleation sites for CaCO_3_ precipitation, (2) pH value, (3) concentration of dissolved inorganic carbon (DIC), and (4) the concentration of free calcium ions in the crystallization solution ^13^. In a MICP process, both the granular medium and the microorganisms can act as nucleation sites for CaCO_3_ precipitation^14^. The remaining three factors influence the saturation state of the solution. The saturation state for CaCO_3_ precipitation is given by^15^:

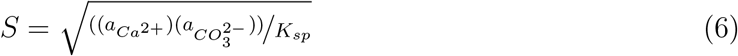

where 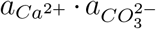 is the ion activity product of calcium and carbonate ions in the solution, and K_sp_ is the solubility product of the nucleating polymorph of CaCO_3_. The system has a driving force towards crystallization once *S* > 1. The solution is then referred to as supersaturated solution. Large S leads to fast spontaneous crystallization, where both crystal nucleation and crystal growth occur. Once crystals nucleate, there is a reduction in solution supersaturation *S*. Below a critical supersaturation level, no new crystal nucleate and crystal growth is responsible for relaxation of the supersaturated system^16^.

In biomineralization processes CaCO_3_ can crystallize in different crystal polymorphs: calcite, vaterite and aragonite as well as hydrated crystalline phases such as monohydrocalcite and ikaite^17^. Calcite is the most common calcium carbonate mineral in nature. It is the most thermodynamically stable form, with the lowest solubility in water^18^. Aragonite is less thermodynamically stable than calcite, but can still be found in nature^19^. Vaterite is the least thermodynamically stable polymorph, and is rarely found in nature. In aqueous solutions, vaterite transforms rapidly into either calcite or aragonite^20^. According to Ostwald’s rule of stages, metastable polymorphs form first and subsequently transform into the thermodynamically more stable polymorph^16^. Additionally, amorphous calcium carbonate (ACC) can be formed during the precipitation process. It rapidly transforms to calcite via vaterite^21^. Furthermore, it has been shown that vaterite preferably crystallizes at high supersaturation levels ^22^.

The schematic precipitation process for MICP is shown in Figure 1. The pH is increased due to urea hydrolysis. Once the solution is supersaturated, spontaneous crystallization occurs. For high supersaturation values, calcite is precipitated via ACC and vaterite, while lower supersaturation levels lead to direct calcite precipitation^23,24^.

**Figure 1:**
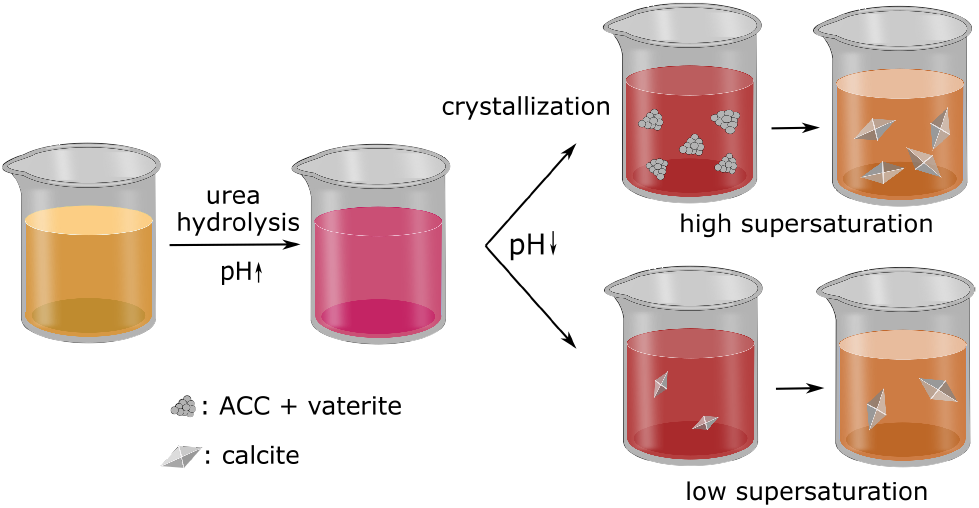
Schematic illustration of the precipitation process: In the first part of the reaction the pH is increasing due to urea hydrolysis. Once the solution is supersaturated crystallization can occur. The crystallization pathway depends on the level of supersaturation. For high supersaturation ACC and vaterite precipitate first, transforming into calcite over time. In the case of low supersaturation, calcite is the favoured nucleating polymorph. Due to crystallization, the pH decreases. The change of the pH value during the crystallization process is indicated by changing colors of pH indicator Phenol Red.

The pH value in a MICP process is controlled by the urea hydrolysis reaction and therefore dependent on the presence of urease. During the ammonium formation the pH increases (Equation 3), and therefore the bicarbonate equilibrium (Equation 4) is shifted. The pH increase leads to an increase of the amount of dissolved inorganic carbon (DIC). Depending on the experimental conditions, two cases can be considered. In a closed system, where there is no gaseous CO_2_ exchange with the atmosphere, the amount of DIC is controlled only by urea hydrolysis. In case of an open system, CO_2_ can also be exchanged with the atmosphere ^25^:

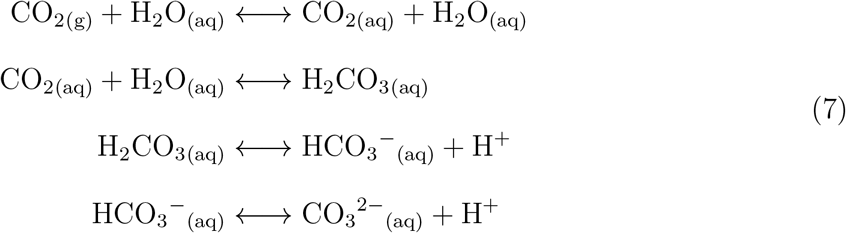

The saturation state of the crystallization solution is also influenced by the calcium concentration. CaCl_2_ is a calcium source most frequently used in MICP^26,27^, however, it is expensive and not environmentally friendly, since Cl^-^ ions react fast to form chloride salts and pollute water. If MICP is used in combination with steel-reinforcement, the presence of chloride ions will promote steel corrosion. The effect of other calcium sources, such as calcium lactate^28,29^ and calcium acetate^29,30^ on properties of the consolidated material have been investigated. Recently, an environmentally friendly and inexpensive alternative based on crushed limestone dissolved with lactic acid was used in a MICP process. In this approach, it is possible to conduct the dissolution process by lactic acid producing bacteria, making both dissolution and precipitation rely on bacterial processes^31^:

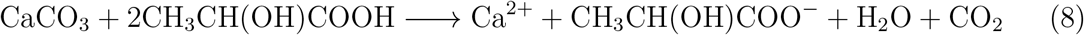

The dissolution process of calcite in aqueous solutions can be characterized in terms of three pH regimes: a regime for low pH values, a transitional regime and a regime for high pH values ^32^. For acidic conditions the dissolution can be in general described with the following equations:

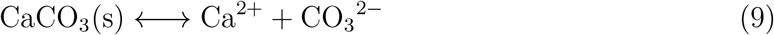

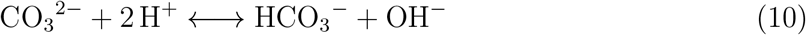

Bicarbonate can react further at very low pH values to gaseous CO_2_:

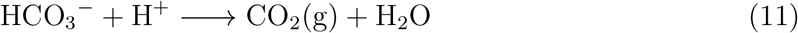

The dissolution reaction consists of several steps such as diffusion of reactant towards the solid surface, adsorption to the surface, migration to an active site on the surface, the dissolution reaction itself on the active site, migration of products away from the active site, desorption of the product, and diffusion of the product into the bulk solution. The slowest of these processes will be the rate limiting process for the dissolution reaction^33^. For pH values below 4 it has been reported that the dissolution is controlled by the diffusion of protons to the calcite surface^34^ and is therefore limited by mass transport. For pH values in the range from pH 4 to pH 5.5 surface reactions are influencing the dissolution, limited by mass transfer.

As a consequence of the dissolution process, calcium and carbonate ions are released into solution. This causes the pH value of the solution to increases (Figure 2).

**Figure 2:**
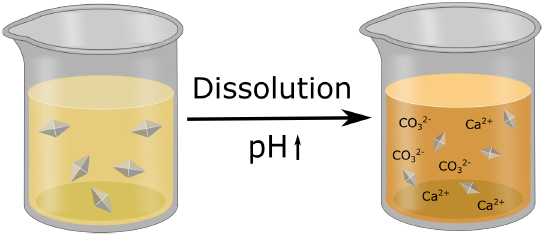
Schematic illustration of the dissolution process: A solution of lactic acid is added to CaCO_3_ crystals. As CaCO_3_ crystals are being dissolved the pH increases over time, leading to a color change of the pH indicator Phenol Red from yellow to orange. As result of the dissolution, Ca^2+^ and CO_3_^2-^ are released into the solution, where CO_3_^2-^ in aqueous solution reacts with bicarbonate and hydroxide ions according to Equation 10.

Previous research in MICP has been focused on macro-scale properties of the consolidated material. Scanning electron microscopy (SEM) is a commonly used technique to investigate characteristics of the precipitated CaCO_3_ minerals (e.g.^35^). Even if properties of the consolidated material, in terms of crystal shape and polymorphism, can be characterized well with imaging methods such as SEM and atomic force microscopy (AFM), these methods fail to provide information about the precipitation process. Microfluidic chips have been developed to observe the formation of CaCO_3_ crystals in real time and *in situ*^36^. Furthermore, nucleation and crystal growth have been investigated with direct optical microscope observation^24^. However, those micro-scale studies do not give information about pH changes during the nucleation and crystallization processes.

In this contribution, we introduce two experimental methods to monitor pH changes in real time and in *situ* in small volumes and at the grain scale. In the first method, the mean pH in small volumes (200 μL) is measured in real time using absorption spectroscopy, and precipitation processes are correlated to pH changes with the help of optical microscopy. The second method allows monitoring of local pH changes at the grain scale with the help of confocal laser scanning microscopy.

Both methods can also be applied to investigate pH changes during the dissolution of CaCO_3_ crystals, for example using lactic acid or acid producing bacteria. The usability of both methods is demonstrated by investigating CaCO_3_ precipitation at high supersaturation induced by *Sporosarcina pasteurii* and CaCO_3_ dissolution with lactic acid.

## Materials and Methods

### Matersials

All chemicals were purchased at SigmaAldrich (Norway) unless otherwise stated. Solutions were prepared with deionized water (DI) and filtered with 0.22 μm polycarbonate syringe-filter before use.

### Crystallization experiments

#### Bacteria culture

For crystallization experiments the urease-producing bacterium *Sporosarcina pasteurii* (Strain DSM33 of *S. pasteurii*) purchased from “Deutsche Sammlung von Mikroorganismen and Zellkulturen” (DSMZ) was used. DSMZ medium 220 with pH of 7.3 supplemented with urea and consisting of 15 g l^-1^ peptone from casein, 5 g l^-1^ peptone from soy meal, 5 g l^-1^ NaCl and 20 gl^-1^ urea was used as growth medium. The culture was inoculated with 1% frozen glycerol stock of strain DSM33. The culture was incubated overnight (17h) at 30 °C with constant shaking. Afterward the culture was transferred to a 50 mL Falcon tube and incubated at 30 ^°^C until further use 2 days after inoculation.

Immediately before use in experiments, the bacterial culture was centrifuged at 5200 *×g* for 8 min and the cells subsequently washed twice with pre-warmed 0.01 M PBS (phosphate buffered saline) to remove the growth medium. The cells were re-suspended and diluted in 0.01 M PBS. Cell concentrations were determined by the optical density measurement of 150 μL dilutions in a 96-well plate at 600 nm (OD_600nm_).

#### Crystallization solution

Dissolved chalk was used as a calcium source^31^. Industrial grade chalk powder (Franzefoss Miljøkalk AS (Norway)), which mainly consists of CaCO_3_, was dissolved with 300mM lactic acid, until the reaction was completed. The solution was left on a shaker for 24 h and afterwards filtered with a 0.22 μm polycarbonate filter to remove any undissolved limestone from the solution. The starting concentration of urea in all crystallization experiments was 0.1 M. To start the crystallization reaction, *S. pasteurii* bacteria suspension was added to the crystallization solution. The ratio of bacteria suspension to the final sample volume was 1:10. As an indication for the bacteria cell concentration of the bacteria suspension the optical density was measured before adding it to the crystallization solution (for the presented experiments: OD_600nm_=1.260).

#### Calcite crystals for dissolution experiments

For dissolution experiments lactic acid (DL-lactic acid 90%) was diluted to a concentration of 10 mM. Calcite crystals were grown directly on Ibidi polymer coverslides (Ibidi GmbH, Gräfelfing, Germany), using the following procedure: The slides were placed in a beaker with CaCl_2_ solution. To initiate the crystallization, Na_2_CO_3_ solution was added and mixed thoroughly. The concentrations of the CaCl2 and Na2CO3 solutions were chosen such that the final concentration of Ca^2+^ and CO_3_^2-^ was 5mM. The reaction was left to proceed for 3 days. Afterwards, the polymer slides were washed with DI water and ethanol before use.

#### Calibration buffers

Calibration buffers in the pH range of 3.1 to 9.5 were used for calibration of the pH sensitive dyes. In the pH range of 3.1 to 6.0 0.5 M MES buffer, and in the pH range of 6.5 to 9.5 0.5 M Tris buffer was used.

#### pH dyes

For global pH monitoring experiments the pH sensitive dye Phenol Red was used. For local pH monitoring experiments the fluorescent pH-sensitive dye, R6G-EDA (N-(rhodamine 6G)-lactam-ethylenediamine)^37,38^ was used in combination with a pH insensitive dye, Sulforhodamine 101 (SR101).

### pH monitoring

#### Global pH monitoring

Absorption spectroscopy was used to monitor pH changes in small volumes (200 μL). Absorption measurements were performed with a spectrophotometer (SpectraMax^®^ i3 Platform). The pH sensitive dye Phenol Red (0.4 μM) was used to make pH changes optically detectable. Phenol Red shows a strong pH dependence at 558 nm and 434 nm, leading to a continuous color change from yellow at low pH to pink at high pH values. The principle of global pH monitoring measurements is shown in Figure 3. Reaction solution and Phenol Red were added to a 96-well plate. After starting the reaction, the well plate was covered with transparent tape to minimize the gas exchange with the environment (Figure 3 (a)). During the reaction the temporal change of the absorption intensity at 558 nm was measured (Figure 3 (b)). In addition, to compensate for scattering due to the crystals in the solution, the background signal at the wavelength of 800 nm was measured, and the background corrected signal was used for pH calculations. A calibration curve was created by measuring the absorption intensity for the buffer solutions with well-known pH values (supplementary information: Figure S1, Figure S2). The background corrected absorption signal of the measurement can then be converted to pH values, and the average pH changes of the sample were detected (supplementary information: Figure S3).

**Figure 3:**
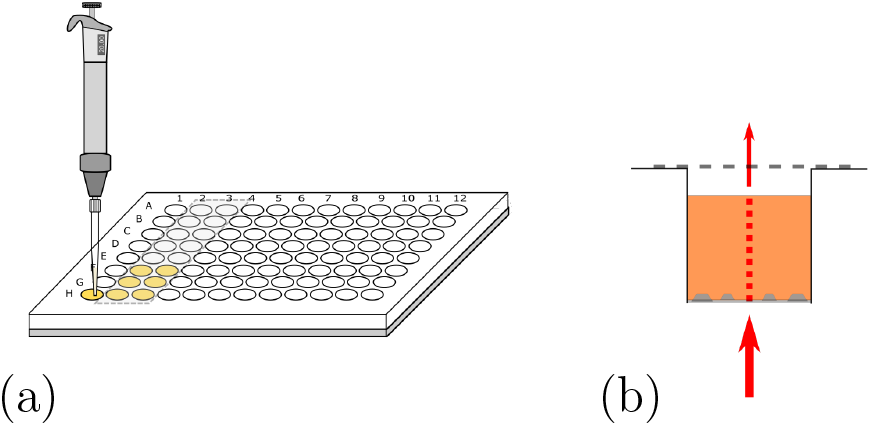
Principle of global pH monitoring for precipitation: (a) *S. pasteurii* dilution is added to the crystallization solution and Phenol Red, pre-filled in the wells of a 96-well plate. To minimize gas exchange with the environment, the wells are sealed with transparent tape after adding the bacteria suspension. (b) Schematic illustration of absorption measurements during crystallization. The amount of absorbed light is measured at various wavelengths.

#### Optical microscopy

Simultaneously to global pH measurements, the precipitation process of a parallel sample in a 96-well plate was investigated with an optical microscope (Motic, AE31E, Objective 20x 0.3NA). Images were taken with a Moticam 5.0.

#### Local pH monitoring

A confocal laser scanning microscope (Leica TCS SP5) (CLSM) was used to investigate local pH evolution on a single crystal level in *situ* and in real time. For that, a fluorescent pH-sensitive dye, R6G-EDA (N-(rhodamine 6G)-lactam-ethylenediamine), was added at a concentration of 0.18 mM to the reaction solution. In addition, a pH insensitive dye, Sulforhodamine 101 (SR101), was added with a final concentration of 15 μM. For measuring local differences of the pH, a flow-cell setup was used (Figure 4 (a)). For the flowcell Ibidi polymer slides and Ibidi sticky-Slide VI 0.4 were used (Ibidi GmbH, Gräfelfing, Germany).

Figure 4 (b) shows a schematic illustration of the local pH monitoring principle. A CLSM is used to scan the area surrounding crystallization or dissolution processes, using a modified technique reported elsewhere^39^. The two fluorescent dyes were excited with an argon laser (excitation wavelength: 514 nm) and a diode pumped solid state laser (excitation wavelength: 561 nm). For spatiotemporal pH measurements the images were scanned sequentially line-

**Figure 4:**
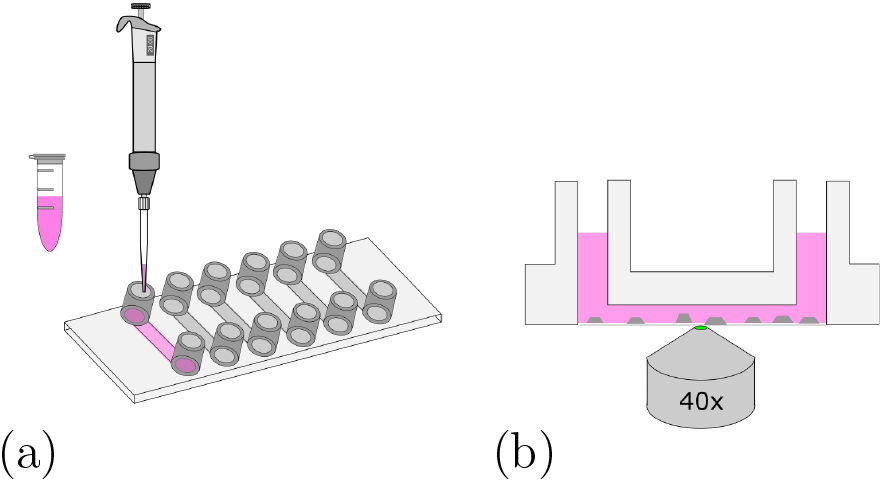
Principle of local pH monitoring for precipitation: (a) Crystallization solution and bacteria dilution are mixed and added to the flow cell. The flow cell is assembled out of an Ibidi Polymer slide and an Ibidi sticky-Slide VI 0.4. (b) Schematic illustration of local pH monitoring during the crystallization with CLSM. The fluorescent dye mixture is excited with two different wavelengths close to crystals to investigate local pH changes.

by-line to avoid cross-talk between the two fluorescent dyes. The fluorescent signal was detected with two Leica HyD detectors in photon counting mode. R6G-EDA and SR101 were detected separately by setting the emission filter to 525 nm to 554 nm and 578 nm to 625 nm, respectively. The scan speed was set to 100 Hz, and image resolution was 512 × 512 pixel. The bit depth was set to 12 bit. All measurements were performed with a pinhole size of 1 Airy. After the measurements, images were processed using a 3× 3 median filter, and the intensity ratio between R6G-EDA and SR101 was calculated pixel-by-pixel. Two standard curves were generated in order to convert the ratio of fluorescent intensities to pH values. For the standard curve, images were taken for 13 pH values in a range from 3.1 to 9.5. The intensity ratios were calculated and fitted with a Sigmoid function:

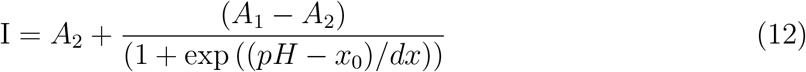

with the fitting parameters *A_1_*: inital value; *A_2_*: final value; *x_0_*: center; *dx:* slope. The curves were used to convert intensity ratios into pH values. For precipitation processes the ratio between SR101 to R6G-EDA was used (further referred to as ratio 1, supplementary information: Figure S4). For dissolution the ratio of fluorescent intensities of R6G-EDA to SR101 was used (in the following referred to as ratio 2, supplementary information: Figure S5). Additionally, a mask was applied to the local pH map images, rendering forming and dissolving crystals gray in the presented pH maps.

### Raman microspectroscopy

For the sample preparation Jack Bean urease (plant derived) was used for catalysing the urea hydrolysis reaction. The precipitation was stopped at different timepoints with ethanol, and the CaCO_3_ precipitate was filtered out of the crystallization solution and dried before further characterization. Raman microspectroscopy was employed to characterize the polymorph phase of the precipitated CaCO_3_ using a Renishaw InVia Reflex Spectrometer equipped with a 532 nm laser and a Leica 50x 0.75NA lens.

## Results

### CaCO_3_ precipitation

For consistency purposes, all experiments presented here were performed with the same batch of *S. pasteurii* (see Materials and Methods). The rate of urea hydrolysis will depend on both, bacteria concentration and in some extent the time-point of the experiment within the growth curve of *S. pasteurii*. Therefore, in the following precipitation experiments, nucleation time-points are specific for the used bacteria cell concentration and growth time.

The precipitation process in a 96-well plate was monitored with optical microscopy to identify different stages of the bacteria-induced precipitation of CaCO_3_. Optical microscopy was initiated approximately 30 sec after the reagents were added (see Materials and Methods). Selected time-points of the crystallization process are shown in Figure 5. Note that the time-labels refer to the start of measurements and are characteristic for the used bacteria concentration.

**Figure 5:**
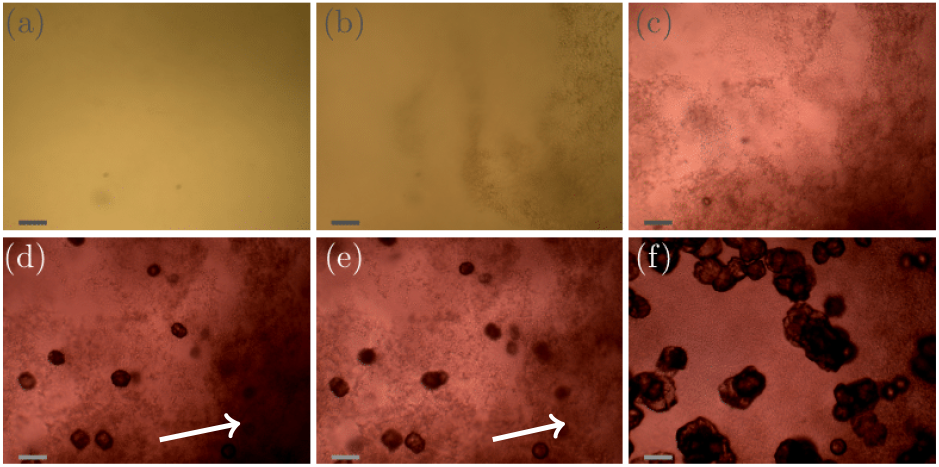
Precipitation time series in a 96-well plate format. Optical microscopy images of precipitation process taken at 6 different time-points: (a) 0 min (= start of measurement), (b) 4 min, (c) 30 min, (d) 50 min, (e) 60 min and (f) 300 min. The precipitation process shows three stages. Stage I (a-b): small particle precipitation appears (2min - 24min). Stage II (c-d): nucleation and initial crystal growth (25 min-50 min), simultaneously the small precipitation from stage I starts to disappear; and Stage III (e-f): the small precipitation from Stage I continues to disappear, and crystal growth is the dominating process (after 50 min). Color changes in the images correspond to a change in the pH value due to the presence of Phenol Red in the sample. The lengths of the scale-bar in the images is 50 μm. The two white arrows indicate an area where the initial precipitation disappears.

Parallel to the optical microscope experiment, the pH evolution of the crystallization solution in multiple samples was monitored using absorption spectroscopy. With the method applied here, the average, i.e. global pH of the solution is measured. This measurements will also serve as a reference for the pH evolution at the grain scale described below.

The pH evolution corresponding to the precipitation process of Figure 5 is shown in Figure 6. Both figures cover the initial 5 h of the reaction. The pH measurements were started approximately 1 min after adding the bacteria cells. For the precipitation process, three distinct stages can be identified. Stage I starts shortly after the start of the reaction and is characterized by the appearance of small particles in the solution. The small particles first float freely in the solution, before settling at the bottom of the well. Figure 5 (a) represents an example for Stage I, 4 min after starting the reaction. The crystallization solution is changing color from yellow to red, indicating a pH increase, as also observed in direct pH measurements (Figure 6). The pH in the starting phase is increasing rapidly up to a value of 8.2.

For the chosen experimental conditions, Stage II starts after approximately 25 min, when the pH of the crystallization solution starts to decrease. As can be seen in Figure 5, this stage is characterized by nucleation and crystal growth occurring simultaneously with dissolution of the precipitates formed in Stage I (see Figure 5b). During Stage II, pH of the solution drops by 0.1 to pH 8.1. Two time-points representing Stage II are shown in Figure 5 c, and d, which illustrate the nucleation, the crystal growth as well as the start of the dissolution of initial precipitated particles (indicated by two white arrows in Figure 5 (d) and (e)).

**Figure 6:**
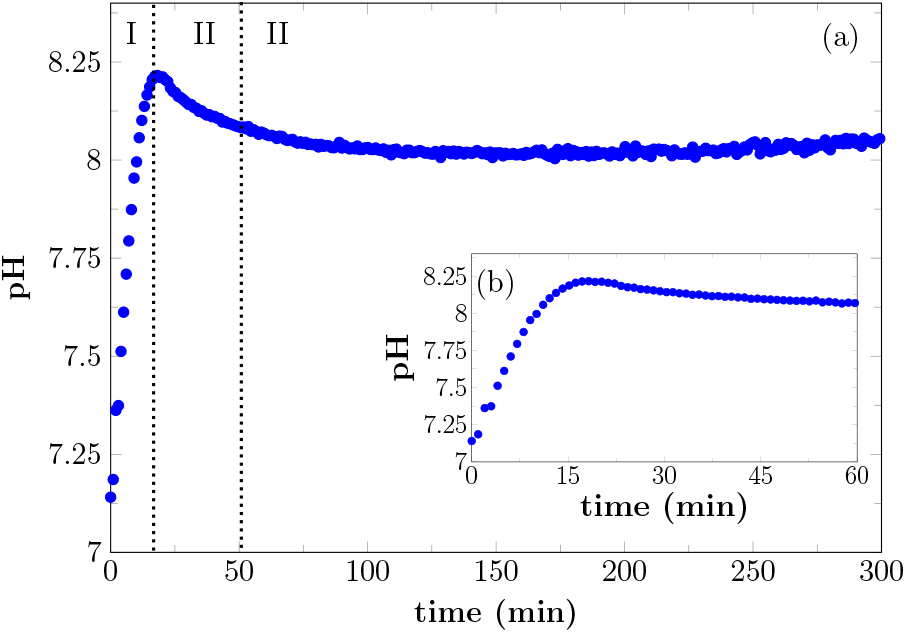
Global pH evolution of *S. pasteurii* induced precipitation (a) over 5 h and (b) the first 60 min of the reaction. The pH increases during the first 17 min of the reaction up to pH 8.2. After a short plateau the pH starts to decrease again after 21 min and reaches a value of 8 after 100 min. A slight increase is seen after approx. 200 min.

In Stage III, which starts in the presented experiment after about 50 min, crystal growth is the dominating process. The initial precipitate continue to dissolve. Correlating this to the global pH evolution measurement (Figure 6) shows, that in this phase, the pH is stable around a value of 8.0. After 200 min the pH of the crystallization solution starts to increase slowly again, which is probably a consequence of slow urea hydrolysis in the absence of free calcium ions, as all calcium is bound in already formed precipitates.

The different polymorphs of the precipitation process have been investigated and identified with Raman microspectroscopy in a separate experiment (Figure 7). In this experiment, plant derived urease was used to catalyse the reaction, and the reaction was stopped by adding ethanol at different time-points (after 1 min, 3min, and 15min). Characteristic Raman peaks for CaCO_3_ can be found at approximately 1085 cm^-1^. It can be seen that the small precipitates that appear initially are consistent with ACC. These quickly transform into vaterite, and a double peak at 1074cm^-1^ and 1085cm^-1^ is observed (see Figure 7). At later time-points (15min), a Raman signal at 1085cm^-1^ and 711 cm^-1^ that is characteristic for calcite crystals is observed. Further appearing Raman peaks at 953 cm^-1^ and 418 cm^-1^ could be caused by intermediate products of the urea hydrolysis reaction as well as by impurities in the formed CaCO_3_. However, the peaks could not be identified with absolute certainty. Despite this, the different polymorph stages during the precipitation process can still be identified as amorphous calcium carbonate (Figure 7 (a)), vaterite (Figure 7 (b)), and calcite (Figure 7 (c)).

**Figure 7:**
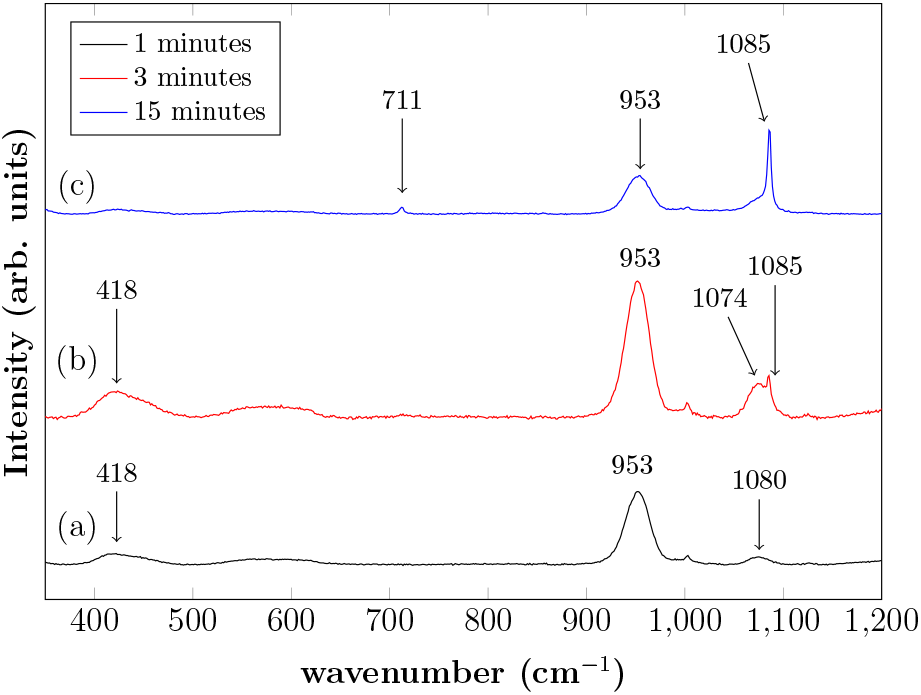
Raman spectra of different precipitation stages. Instead of urease produced by bacteria, plant derived urease was used for those experiments. Precipitation was characterized after (a) 1 min, (b) 3 min and (c) 15 min reaction times. Characteristic Raman peaks for CaCO_3_ can be found at approximately 1085 cm^-1^. (a) A broad peak can be identified at 1080 cm^-1^, which corresponds to amorphous calcium carbonate. With increasing time the amorphous calcium carbonate transforms into a vaterite phase (b) (double peak at 1074 cm^-1^ and 1085 cm^-1^). After 15 min a characteristic calcite peak can be identified at 1085 cm^-1^. In addition, a peak at 711 cm^-1^ can be observed which is characteristic for calcite. Furthermore, peaks at 953 cm^-1^ (a,b,c) and 418 cm^-1^ (a,b) could be observed.

In addition to investigating the pH changes during the precipitation process at the global scale in small volumes, pH evolution was also monitored at the grain scale using CLSM microscopy. 2D local pH changes during the precipitation process close to the precipitating crystals could be documented, using the same concentration of urea, calcium and bacterial cells.

A representative example of spatial and temporal pH mapping is shown in Figure 8. The pH evolution at the scale of individual crystals is shown in Figure 8 (a-h), with the corresponding brightfield images in Figure 8 (i-p). Data acquisition started approximately 2 min after mixing the reactants (Materials and Methods). For easier identification of local pH differences, two selected areas in each of the images are highlighted and magnified. The upper area corresponds to a location in the sample where a calcite crystal nucleates after 12 min. The lower area shows pH evolution in a region in which no calcite nucleation occurs during the course of the experiment. Local pH evolution is monitored for the first 90 min of the reaction.

**Figure 8:**
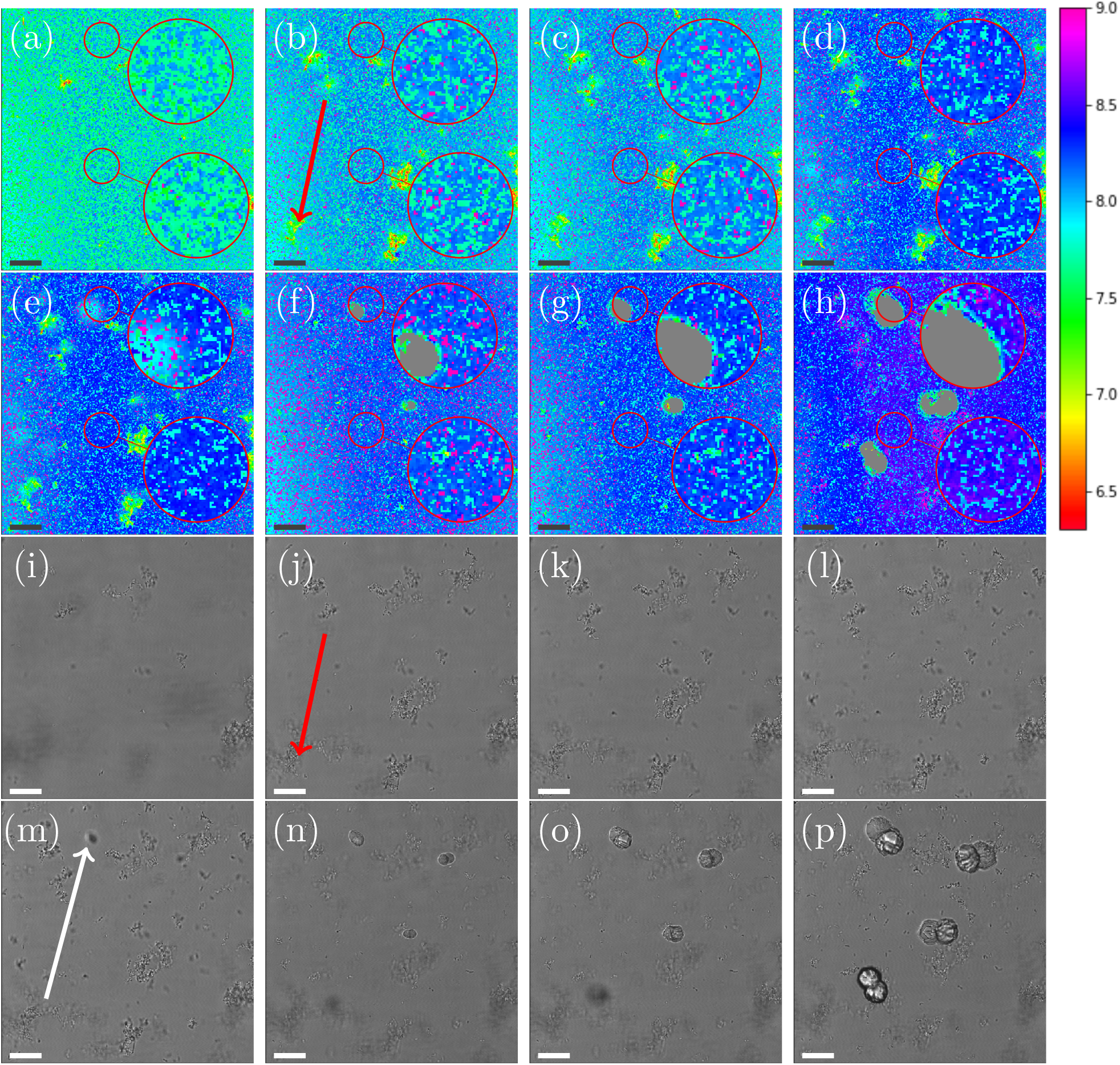
Local pH monitoring of the precipitation process. (a)-(h) Local pH differences during the process are shown with a colormap for different timepoints (2 min, 4 min, 6 min, 12 min, 14 min, 17 min, 22 min, and 62 min) of the reaction. (i)-(p) are the corresponding optical images to the local pH map. During the first 14 min of the reaction, the pH increases, and in areas of precipitation, the pH is lower compared to the rest of the sample. The first crystallization events are observed after 14 min. The pH around the crystals is lower compared to areas further away from the crystals. For clarification, two areas of the images are magnified. The upper zoom magnifies an area close to a nucleation point, while the lower zoom shows an area without crystallization. The pH is color-coded in the images (a)-(f) and the corresponding pH value can be identified with the help of the color-bar on the side. The pH is color-coded and the corresponding pH value can be identified with the color-bar on the side. The shown scale-bar in all images is 25 μm.

As in the global precipitation experiments, three distinct stages of precipitation can be observed. Note that the timescale is not exactly the same as in the experiments at the global scale (Figure 5). This is due to differences between the experimental setups that affect factors such as gas exchange with the environment, local variations in temperature, and different crystallization conditions.

Figure 8 shows the spatial and temporal pH evolution during the precipitation process. In Stage I, during the first 12 min, the color changes from light green to blue, corresponding to a continuous pH change from around 7.7 to 8.2. In good agreement with the global-scale experiments, small particles of ACC and vaterite precipitate and float in the liquid. In areas where ACC and vaterite appears, the local pH decreases (to around pH 7.0) compared with the rest of the sample. An example of this is indicated by a red arrow in Figure 8 (b) and (j). The first calcite crystal can be observed after 14 min, which marks the beginning of Stage II of the crystallization process. Calcite crystals first appear as dark spots in the optical image (Figure 8 (m); marked with an arrow). Simultaneously, it can be observed (Figure 8 (e)) that the pH around the growing crystal is lowered by approximately 0.5 compared with the rest of the sample (pH 7.9).

The crystallization process is shown in more detail in Figure 9. The highlighted and magnified part emphasizes the area where crystallization occurs. Here, it shows that the dark spot on the left side starts to appear after 13 min in the brightfield image. Simultaneously, the spatial pH mapping shows a pH drop in the same area. This can be clearer seen in Figure 9 (c) and (d). A color change to cyan can be observed in the area where the crystal is appearing. This corresponds to a pH drop from 8.2 to 7.9. During crystal growth (Figure 8 (f)-(h)), a color change to green/cyan can be observed in the area close to the growing crystal. This is equivalent to a pH in a range between 7.5 to 8.0. The pH of the rest of the sample is above 8.2 (blue color Figure 8 (e)-(h)). The crystal growth corresponds to the Stage III of the precipitation process. Additionally, it can be observed that the ACC and vaterite formed in Stage I and visible as small particles in the brightfield images, as well as local pH changes in the pH mapping, is starting to dissolve, and the nucleated crystals increase in size. The dissolution of vaterite can be observed after crystallization has started. This also coincides with a more uniform pH map in areas without growing crystals (see Figure 8 (h); lower magnified area).

**Figure 9:**
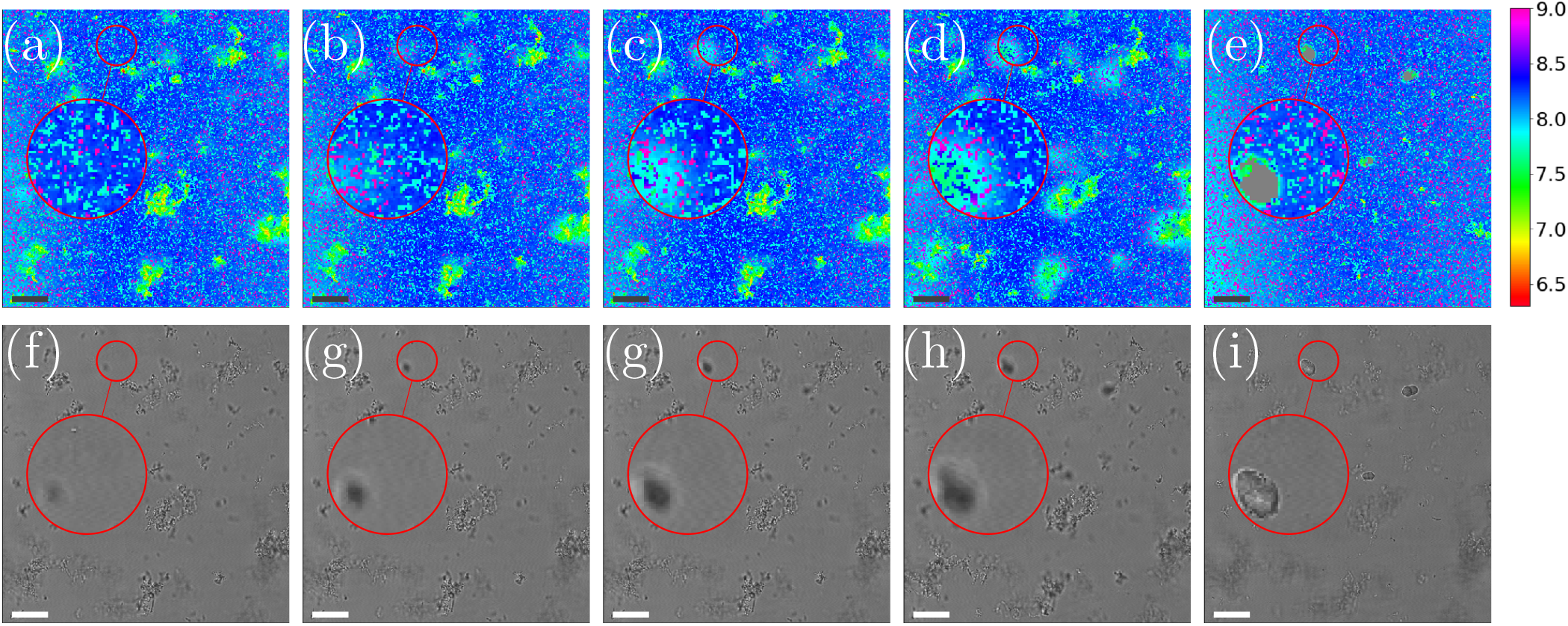
Spatial pH map during the crystallization process (a)-(e) and corresponding bright-field images (f)-(i). The shown time-points are 12min, 13min, 14min, 15min, 16min after the start of the experiment. Calcite growth starts out of focus after 11 min. The crystal increases in size and appears in the focus after 14 min. Spatial pH distribution shows lower pH during the crystallization (b,c,d). The pH is color-coded and can be identified with the color-bar on the side. The scale-bar in all images is 25 μm.

### Dissolution of CaCO_3_ crystals

Spatial and temporal evolution of the pH at the scale of single CaCO_3_ crystals was investigated in dissolution experiments. Our goal is to employ bacteria to produce acid for the dissolution of CaCO_3_, and therefore a method was developed to study the local pH variations during calcite dissolution. For the presented experiments calcite crystals were grown on a polymer microscope slide (see Materials and Methods) and then dissolved using 10 mM aqueous solution of lactic acid. Experiments were conducted in a flow cell as shown in Figure 4 (a) under stagnant (no flow) conditions. Addition of the acid leads to slow dissolution of the calcite crystals as observed by decreasing crystal size (Figure 10) and to changes in the solution pH (Figure 11).

**Figure 10:**
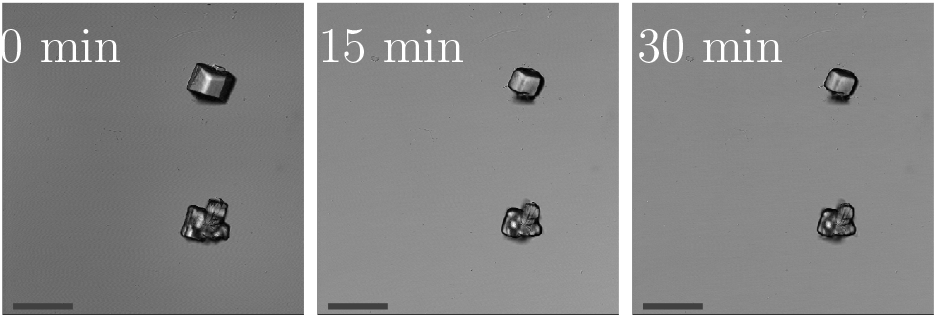
Dissolution of calcite crystals with lactic acid right after adding the acid to the flow cell (=0min), after 15min and 30min reaction time. The scale-bar corresponds to 75 μm.

**Figure 11:**
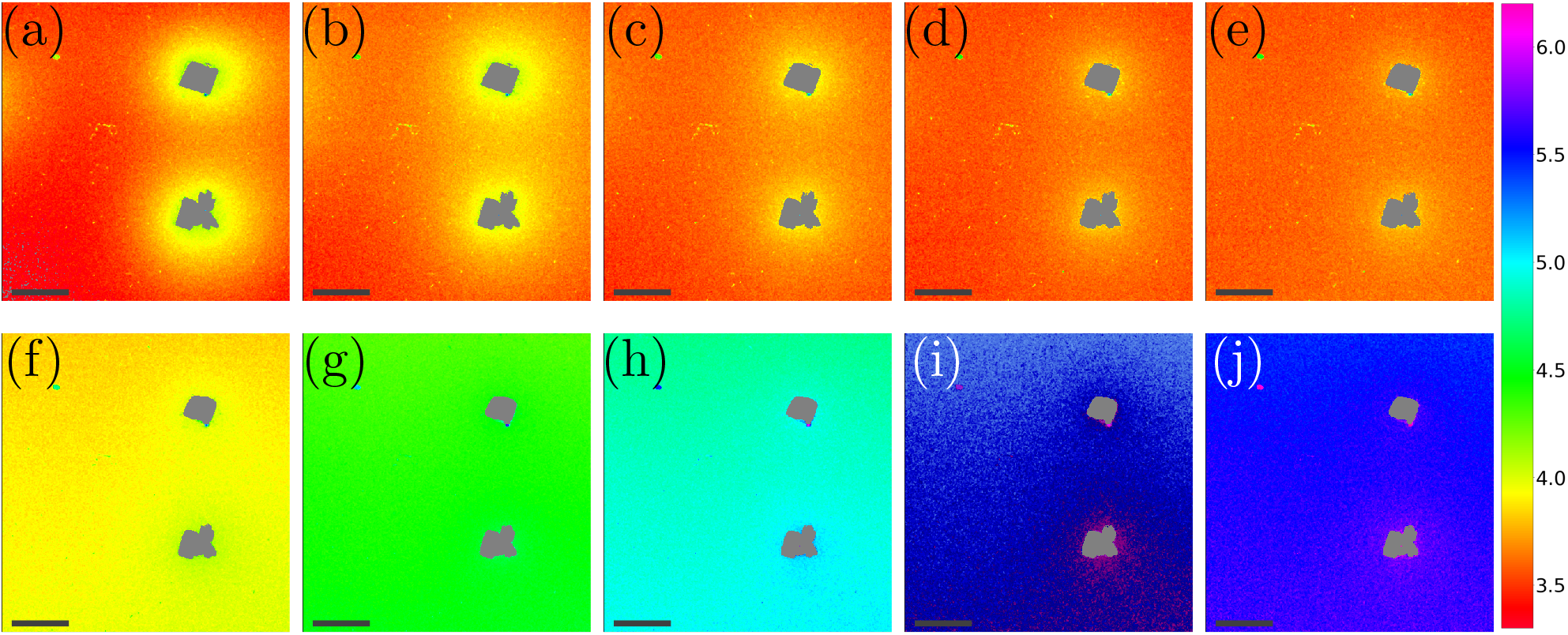
Local pH monitoring of the dissolution process.(a)-(e) shows time-points at the start of the measurement (=0min), after 1min, 2 min, 3 min and 5 min of the measurement. Also later time-points of the dissolution process are shown, i.e. (f) 10min, (g) 15min, (h) 20 min, (i) 25 min and (j) 30 min. The pH is color-coded and the corresponding pH value can be identified with the help of the color-bar on the side. The scale-bar corresponds to 75 μm.

The dissolution process of calcite crystals was investigated with CLSM using the same approach as used to monitor local pH changes in bacteria induced precipitation described above. Measurements are started approximately 30 sec after the addition of the acid. The results of the microscale pH mapping are shown in Figure 11.

Shortly after the acid is introduced into the flow cell, clearly observable local pH gradients are established in the proximity of dissolving crystals. Regions with increased pH can be recognized by the bright yellow color in Figure 11 (a). The pH value close to the crystal is around 4.1, while the pH further away from the crystals is around 3.4. During the first 5 min of the experiment, the area with higher pH is decreasing, while the pH of the entire sample increases.

During the dissolution process, two phases can be observed. In the starting phase of the dissolution, the main observed process is the pH change in close proximity of the calcite crystal. The pH close to the crystal is decreasing, while the pH of points distant to the crystals is increasing. Once a uniform pH in the sample is established (after approximately 5 min), the main observed process in local pH monitoring experiments is the increase of the pH of the entire sample.

The pH profiles have been plotted along the dashed line shown in Figure 12 (a) for 7 different time-points. Local pH differences are largest at the beginning of the dissolution process (0min). The pH value close to the crystal is around 4.1, while the pH further away is around 3.4. During the initial 0 min to 5 min, the local differences in pH become smaller, until after 5 min the pH is nearly uniform in the sample. A higher pH near the dissolving crystals can only be detected within a distance of 40 μm from the crystal surfaces.

## Discussion

### CaCO_3_ precipitation

During CaCO_3_ precipitation, we observed the formation of different CaCO_3_ polymorphs (Figure 5, Figure 8). In the initial stage of the reaction, ACC and vaterite phases were observed, while at later time-points of the experiment calcite was the dominating polymorph (Figure 7). It has been previously reported that during CaCO_3_ precipitation at high supersaturation conditions, hydrated ACC is formed first^21^. This phase subsequently transforms into vaterite. The metastable vaterite dissolves again once the supersaturation with respect to vaterite approaches *S* =1 and according to Ostwald’s rule of stages, reprecipitates as the most stable CaCO_3_ polymorph, that is calcite. This has also been reported for MICP processes^23^. The precipitation and dissolution of small CaCO_3_ particles in MICP has previously been observed in microscale experiments^24^. In our experiments, the previously reported precipitation pathway could be confirmed. A further confirmation of the different polymorphs during the MICP precipitation has been achieved using Raman microspectroscopy (Figure 7).

The presented global pH monitoring method provides information about the average pH evolution of the crystallization solution. The expected pH evolution is a rapid pH increase due to urea hydrolysis followed by a pH decrease due to crystallization of CaCO_3_ (see Equation 5), which was confirmed by our measurements (Figure 6). This also agrees with previously reported pH evolution’s in bulk experiments. In the work of Dupraz *et al.*^40^, the pH evolution in larger volumes (30 mL and 60 mL) was investigated with and without gas exchange with the environment, showing the influence of gas exchange on the pH evolution. In systems with limited gas exchange, the pH increase and carbonate precipitation was faster than in systems with gas exchange with the environment (see Equation 7). In addition, the pH value before nucleation was reported to be higher for closed systems. This is due to limitation of CO_2_ during the hydrolysis reaction. CO_2_ exchange with the environments is more pronounced when the surface to volume ratio of the sample is high, which is the case in our global pH monitoring experiments. This explains why the global pH values we detect are low compared with previously reported MICP experiments^10,40^.

In our observations of local pH on the grain scale, we found that both the appearance of the initial precipitation (ACC and vaterite), as well as the crystallization of calcite, gave rise to local decrease in pH. The pH decrease due to ACC and vaterite precipitation could not be detected in the global pH measurements, because its average effect was smaller than the global pH increase due to urea hydrolysis.

The precipitation processes in global and local pH monitoring experiments show the same trend, however crystallization occurs at different time-points, despite using of the same bacteria cell concentrations. This can be explained by several experimental factors. Perhaps the most important is that the flow cell (Figure 4 (a)) can be considered a closed system, because only a very small fraction of the solution is in equilibrium with the atmosphere. In the global pH monitoring experiments, gas exchange was limited, however, gas exchange can not be neglected for those measurements. The closed conditions of the flow cell experiments lead to a faster pH increase and as a consequence faster precipitation of CaCO_3_.

Furthermore, the influence of the surface morphology of the substrate on precipitation and morphology of precipitated crystals in MICP has been reported earlier^41^. This shows that precipitation depends on the surface chemistry of the substrate, and can contribute to the different time-points at which precipitation starts in global and local pH monitoring experiments.

### Dissolution of CaCO_3_ crystals

In our dissolution experiments, we observe a slow dissolution of calcite crystals (Figure 10). H^+^ ions are consumed in the dissolution process (Equation 10), and the resulting diffusion of protons from the bulk solution towards the dissolving surfaces can be seen as a pH gradient between the surface and the bulk solution. After a reaction time of 5 min, a pH profile with nearly constant shape is reached (Figure 11).

At low pH values, it is expected that the bicarbonate from the dissolution reaction further reacts to produce CO_2_. This could not be observed in the presented experiments due to fast diffusion of the reaction products and low concentration of dissolved carbon.

As shown in Figure 12, the pH close to the crystal is observed to decrease after a very fast increase from bulk pH conditions, during the first 5 min of the experiment. This somewhat counter-intuitive observation can probably be explained by an initial fast dissolution rate of the calcite crystal, caused by the large availability of active reaction sites on the fresh calcite surface^42^. As the density of reactive sites decreases and the dissolution rate slows down, the diffusion of H^+^ and reaction products causes a temporary drop in local pH. This previously unobserved dynamics may have implications for biogeochemical reactions in the vicinity of dissolving calcite surfaces.

**Figure 12:**
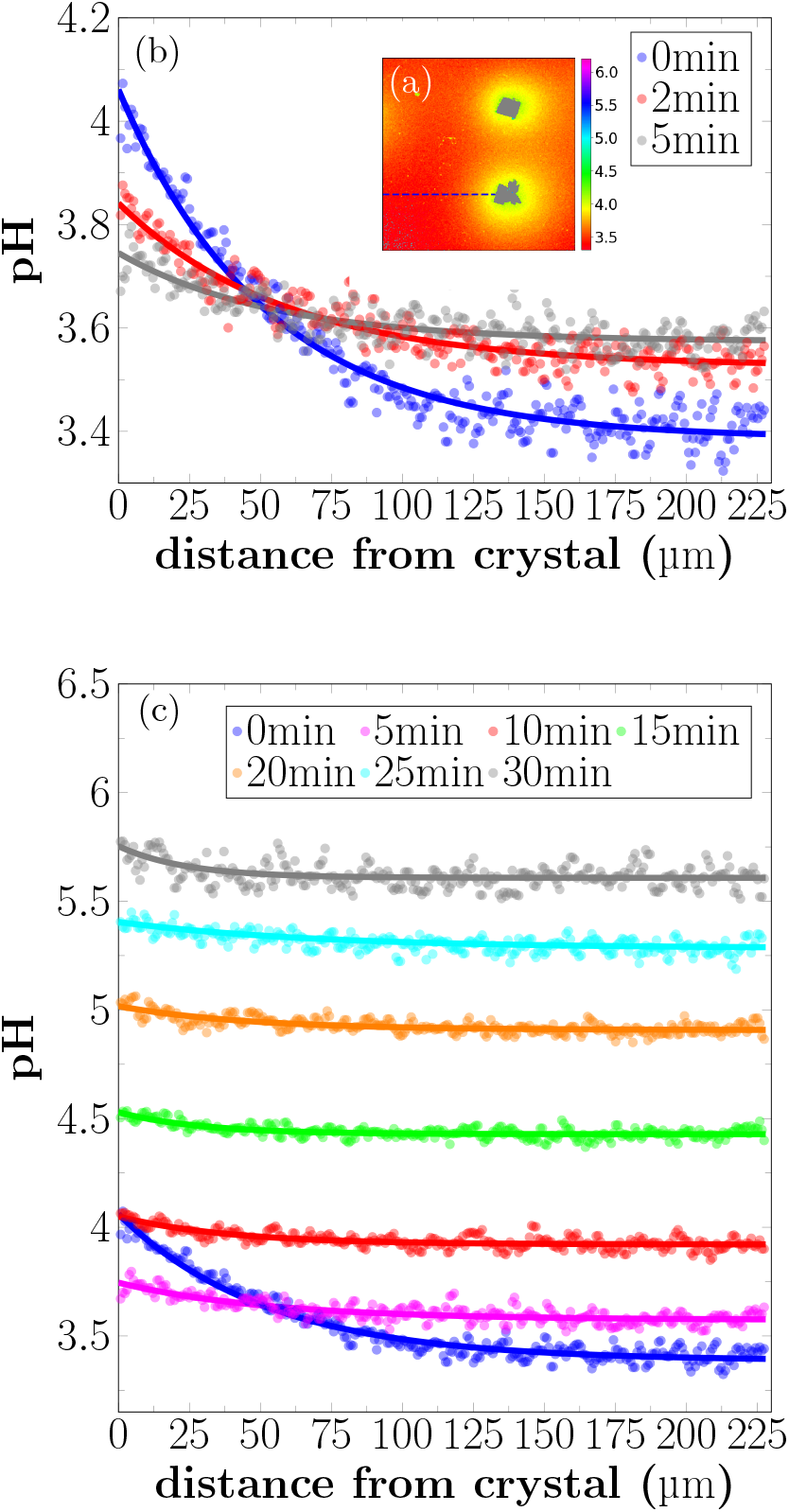
pH profile close to a dissolving crystal for different time-points of dissolution. (a) Example image of local pH distribution during dissolution process. (b) Intensity profiles are extracted from the images along the dashed line. Intensity profiles are given 0 min, 2 min, and 5 min of the reaction. Differences in the local pH are first limited close to the crystal. With ongoing reaction time, local differences spread out and the pH value of the whole sample is increasing. (c) pH profiles for longer reaction times. Over time, local pH differences decrease, and the pH of the whole sample increases.

## Conclusions

We have presented two novel methods for monitoring the pH evolution of MICP processes at the micro-scale. The global pH evolution in small volumes was measured with absorption spectroscopy, where the pH is detected due to the presence of the pH sensitive dye Phenol Red in the sample. Furthermore, a method was introduced for monitoring local pH changes using CLSM with one pH sensitive and one pH insensitive fluorescent dye.

With the presented methods we were able to achieve detailed, *in situ* pH monitoring, which allows us to learn more about the reactions during MICP. This allows us to gain more control when designing and optimising the process on the micro-scale.

Local pH monitoring revealed details about local pH changes, which have not been observed in bulk measurements. We were able to demonstrate that the appearance of metastable phases during the initial pH increase causes a local pH decrease. Furthermore, for dissolution experiments local and temporal pH variations around the dissolving calcite crystals have been observed, which may have implications for other biochemical reactions taking place in close proximity of the dissolving crystals.

The methods introduced in this work allow the detailed, micro-scale evaluation of MICP processes and represent a valuable tool for improving material properties in MICP-based materials, with prospects for becoming more sustainable alternatives for conventional concrete in a wider range of applications in the construction industry.

## Acknowledgement

This work was supported by the Research Council of Norway under project 269084. Additional support was provided by Norwegian Micro- and Nano-Fabrication Facility, NorFab under Research Council of Norway project 245963/F50. The authors thank Simone Balzer Le and Sidsel Markussen from SINTEF Industry, Trondheim, Norway for providing the bacteria cultures for this work.

## Supporting Information Available

The following figures are available in the supplementary information:

- Phenolredabsorption.pdf: Figure S1: Absorption spectra of pH sensitive dye for pH values between 3.2 and 9.5
- Phenolredcalibrationcurve.pdf: Figure S2: Calibration curve for global pH monitoring experiments.
- Absorptionsignal.pdf: Figure S3: Signal processing during global pH monitoring experiments
- localpHratio1.pdf: Figure S4: Calibration curve for ratio 1 for local pH monitoring experiments
- localpHratio2.pdf: Figure S5: Calibration curve for ratio 2 for local pH monitoring experimentsGraphical TOC Entry

## References

(1) Le Quéré, C.; Moriarty, R.; Andrew, R. M.; Canadell, J. G.; Sitch, S.; Korsbakken, J. I.; Friedlingstein, P.; Peters, G. P.; Andres, R. J.; Boden, T. A., et al. Global carbon budget 2015. Earth System Science Data 2015, 7, 349–396.

(2) Agency, I. E. Cement Technology Roadmap: Carbon Emissions Reductions up to 2050; 2009; p 36.

(3) Naqi, A.; Jang, J. G. Recent progress in green cement technology utilizing low-carbon emission fuels and raw materials: A review. Sustainability 2019, 11, 537.

(4) Bosoaga, A.; Masek, O.; Oakey, J. E. CO2 capture technologies for cement industry. Energy procedia 2009, 1, 133–140.

(5) Vatopoulos, K.; Tzimas, E. Assessment of CO2 capture technologies in cement manufacturing process. Journal of cleaner production 2012, 32, 251–261.

(6) Achal, V.; Mukherjee, A. A review of microbial precipitation for sustainable construction. Construction and Building Materials 2015, 93, 1224–1235.

(7) De Muynck, W.; De Belie, N.; Verstraete, W. Microbial carbonate precipitation in construction materials: a review. Ecological Engineering 2010, 36, 118–136.

(8) Stabnikov, V.; Naeimi, M.; Ivanov, V.; Chu, J. Formation of water-impermeable crust on sand surface using biocement. Cement and Concrete Research 2011, 41, 1143–1149.

(9) Wong, L. S. Microbial cementation of ureolytic bacteria from the genus Bacillus: a review of the bacterial application on cement-based materials for cleaner production. Journal of Cleaner Production 2015, 93, 5–17.

(10) Stocks-Fischer, S.; Galinat, J. K.; Bang, S. S. Microbiological precipitation of CaCO3. Soil Biology and Biochemistry 1999, 31, 1563–1571.

(11) Achal, V.; Mukherjee, A.; Basu, P. C.; Reddy, M. S. Strain improvement of Sporosarcina pasteurii for enhanced urease and calcite production. Journal of Industrial Microbiology and Biotechnology 2009, 36, 981–988.

(12) Sarda, D.; Choonia, H. S.; Sarode, D. D.; Lele, S. S. Biocalcification by Bacillus pasteurii urease: A novel application. Journal of Industrial Microbiology and Biotechnology 2009, 36, 1111–1115.

(13) Hammes, F.; Verstraete, W. Key roles of pH and calcium metabolism in microbial carbonate precipitation. Reviews in Environmental Science and Biotechnology 2002, *1*, 3–7.

(14) Ghosh, T.; Bhaduri, S.; Montemagno, C.; Kumar, A. Sporosarcina pasteurii can form nanoscale calcium carbonate crystals on cell surface. PloS one 2019, *14*.

(15) Morse, J. W. The kinetics of calcium carbonate dissolution and precipitation. Carbonates: Mineralogy and Chemistry, Reviews in Mineralogy 1983, 11, 227–264.

(16) Mullin, J. Crystallization; Chemical, Petrochemical & Process; Elsevier Science, 2001.

(17) Cölfen, H.; Fricke, M.; Harry, S.; Imai, H.; Kniep, R.; Sato, K.; Sewell, S.; Simon, P.; Volkmer, D.; Wright, D. Biomineralization I: Crystallization and Self-Organization Process; Springer Science & Business Media, 2007; Vol. 1.

(18) Boulos, R. A.; Zhang, F.; Tjandra, E. S.; Martin, A. D.; Spagnoli, D.; Raston, C. L. Spinning up the polymorphs of calcium carbonate. Scientific Reports 2014, 4, 1–6.

(19) Weiner, S. Microarchaeology: beyond the visible archaeological record; Cambridge University Press, 2010.

(20) Ni, M.; Ratner, B. D. Differentiating calcium carbonate polymorphs by surface analysis techniques-An XPS and TOF-SIMS study. Surface and Interface Analysis 2008, 40, 1356–1361.

(21) Rodriguez-Blanco, J. D.; Shaw, S.; Benning, L. G. The kinetics and mechanisms of amorphous calcium carbonate (ACC) crystallization to calcite, via vaterite. Nanoscale 2011, 3, 265–271.

(22) Han, Y. S.; Hadiko, G.; Fuji, M.; Takahashi, M. Crystallization and transformation of vaterite at controlled pH. Journal of crystal growth 2006, 289, 269–274.

(23) van Paassen, L. Biogrout, ground improvement by microbial induced carbonate precipitation. Ph.D. thesis, 2009.

(24) Zhang, W.; Ju, Y.; Zong, Y.; Qi, H.; Zhao, K. In Situ Real-Time Study on Dynamics of Microbially Induced Calcium Carbonate Precipitation at a Single-Cell Level. Environmental Science & Technology 2018, 52, 9266–9276.

(25) Stumm, W.; Morgan, J. Aquatic chemistry: an introduction emphasizing chemical equilibria in natural waters; Environmental Science and Technology: A Wiley-Interscience Series of Texts and Monographs; Wiley, 1981.

(26) DeJong, J. T.; Fritzges, M. B.; Nüsslein, K. Microbially Induced Cementation to Control Sand Response to Undrained Shear. Journal of Geotechnical and Geoenvironmental Engineering 2006, 132, 1381–1392.

(27) Burbank, M.; Weaver, T.; Lewis, R.; Williams, T.; Williams, B.; Crawford, R. Geotechnical tests of sands following bioinduced calcite precipitation catalyzed by indigenous bacteria. Journal of Geotechnical and Geoenvironmental Engineering 2013, 139, 928936.

(28) Xu, J.; Du, Y.; Jiang, Z.; She, A. Effects of Calcium Source on Biochemical Properties of Microbial CaCO3 Precipitation. Frontiers in Microbiology 2015, 6, 1366.

(29) Gorospe, C. M.; Han, S. H.; Kim, S. G.; Park, J. Y.; Kang, C. H.; Jeong, J. H.; So, J. S. Effects of different calcium salts on calcium carbonate crystal formation by Sporosarcina pasteurii KCTC 3558. Biotechnology and Bioprocess Engineering 2013, 18, 903–908.

(30) Zhang, Y.; Guo, H. X.; Cheng, X. H. Role of calcium sources in the strength and microstructure of microbial mortar. Construction and Building Materials 2015, 77, 160–167.

(31) Røyne, A.; Phua, Y. J.; Le, S. B.; Eikjeland, I. G.; Josefsen, K. D.; Markussen, S.; Myhr, A.; Throne-Holst, H.; Sikorski, P.; Wentzel, A. Towards a low CO2 emission building material employing bacterial metabolism (1/2): The bacterial system and prototype production. PloS one 2019, 14.

(32) Sjöberg, E. L.; Rickard, D. T. Temperature dependence of calcite dissolution kinetics between 1 and 62C at pH 2.7 to 8.4 in aqueous solutions. Geochimica et Cosmochimica Acta 1984, 48, 485–493.

(33) Morse, J. W.; Arvidson, R. S. The dissolution kinetics of major sedimentary carbonate minerals. Earth-Science Reviews 2002, 58, 51–84.

(34) Sjöoberg, E. L.; Rickard, D. T. Calcite dissolution kinetics: Surface speciation and the origin of the variable pH dependence. Chemical Geology 1984, 42, 119–136.

(35) DeJong, J. T.; Mortensen, B. M.; Martinez, B. C.; Nelson, D. C. Bio-mediated soil improvement. Ecological Engineering 2010, 36, 197–210.

(36) Wang, Y.; Soga, K.; Dejong, J. T.; Kabla, A. J. A microfluidic chip and its use in characterising the particle-scale behaviour of microbial-induced calcium carbonate precipitation (MICP). Géotechnique 2019, 69, 1086–1094.

(37) Li, Z.; Wu, S.; Han, J.; Han, S. Imaging of intracellular acidic compartments with a sensitive rhodamine based fluorogenic pH sensor. Analyst 2011, 136, 3698–3706.

(38) Wu, J.-S.; Hwang, I.-C.; Kim, K. S.; Kim, J. S. Rhodamine-based Hg2+-selective chemodosimeter in aqueous solution: fluorescent OFF-ON. Organic Letters 2007, *9*, 907–910.

(39) Bjørnøy, S. H.; Mandaric, S.; Bassett, D. C.; Åslund, A. K.; Ucar, S.; Andreassen, J.-P.; Strand, B. L.; Sikorski, P. Gelling kinetics and in situ mineralization of alginate hydrogels: A correlative spatiotemporal characterization toolbox. Acta biomaterialia 2016, 44, 243–253.

(40) Dupraz, S.; Parmentier, M.; Ménez, B.; Guyot, F. Experimental and numerical modeling of bacterially induced pH increase and calcite precipitation in saline aquifers. Chemical Geology 2009, 265, 44–53.

(41) Rodriguez-Navarro, C.; Jroundi, F.; Schiro, M.; Ruiz-Agudo, E.; Gonzélez-Munoz, M. T. Influence of substrate mineralogy on bacterial mineralization of calcium carbonate: implications for stone conservation. Appl. Environ. Microbiol. 2012, 78, 4017–4029.

(42) Bibi, I.; Arvidson, R. S.; Fischer, C.; Lüttge, A. Temporal evolution of calcite surface dissolution kinetics. Minerals 2018, 8, 256.

