## Supplementary document for "Microbial-induced calcium carbonate precipitation: An experimental toolbox for in situ and real-time investigation of micro-scale pH evolution"

**Abstract**

In this contribution we present two experimental methods to monitor pH changes in small volumes for microbial-induced calcium carbonate precipitation (MICP). The global pH monitoring method monitors pH changes in small volumes in real time using absorption spectroscopy. The monitored pH changes have been correlated with precipitation processes in the sample with help of optical microscopy. In addition, local pH changes have been monitored on a grain scale with confocal laser scanning microscopy (CLSM) in real time and *in situ*. The supplementary information contains calibration curves and signal processing information for the presented methods.

$$\mathrm{I} = A_2 + \frac{(A_1 - A_2)}{(1 + \exp\left((pH - x_0)/dx\right))} \tag{1}$$

with the fitting parameters $A_1$: inital value; $A_2$: final value; $x_0$: center; $dx$: slope. The curves were used to convert intensity ratios into pH values. For precipitation processes the ratio between SR101 to R6G-EDA was used (further referred to as ratio 1, Figure S4). For dissolution the ratio of fluorescent intensities of R6G-EDA to SR101 was used (in the following referred to as ratio 2, Figure S5).

### Supplementary Material

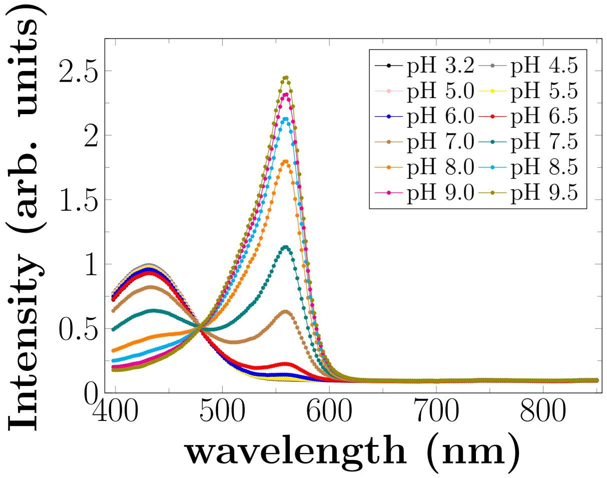


Figure S1: Absorption spectra of pH sensitive dye Phenol Red. The pH sensitive dye Phenol Red shows two pH dependent absorption maxima at 558 nm and 434 nm. The background of measurements can be detected at wavelengths in the range of 650 nm to 850 nm since in this areas, there is no signal from the dye itself.

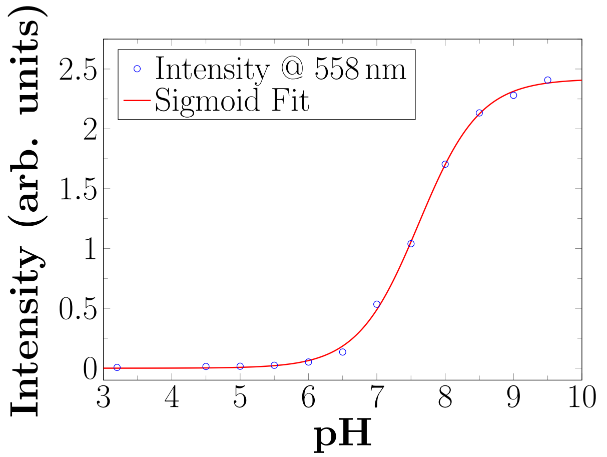


Figure S2: Calibration curve for global pH measurements. The intensity values of the background corrected absorption maxima at 558 nm in dependence of the pH are fitted with a sigmoid function.

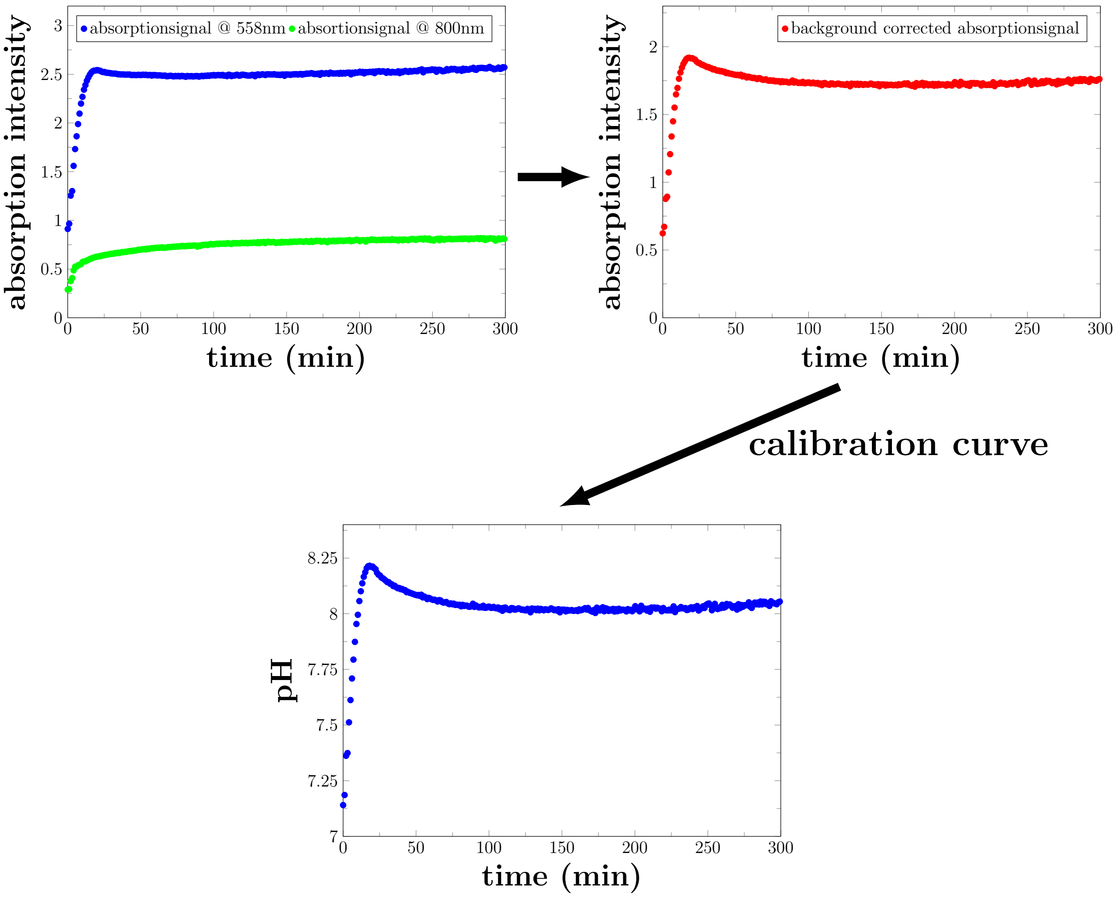


Figure S3: Signal processing during global pH monitoring measurements: Absorption intensity at 558 nm and background signal at 800 nm are recorded over time. Substracting the two signals results in the background corrected signal. The background corrected signal can be converted to pH values with the help of the calibration curve (Figure S2).

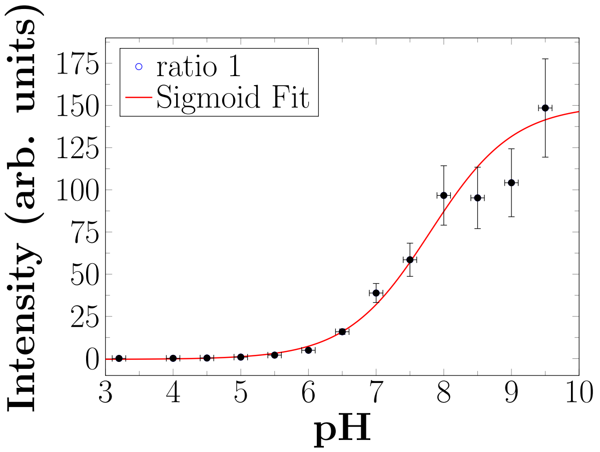


Figure S4: Calibration curve (ratio 1) for local pH monitoring in precipitation experiments. The calibration values were fitted with a sigmoid function.

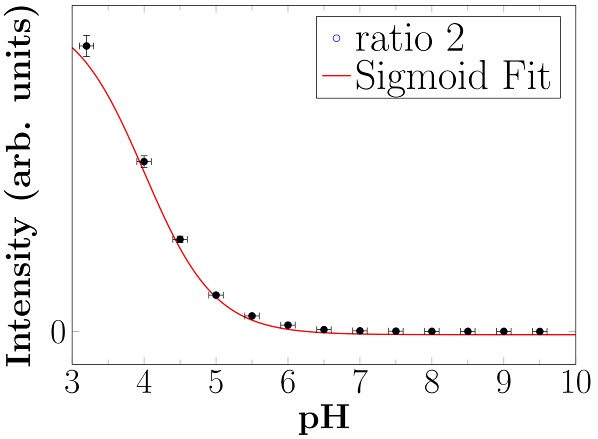


Figure S5: Calibration curve (ratio2) for local pH monitoring in dissolution experiments. The calibration values were fitted with a sigmoid function.
